# Proteolytically Coordinated Activation of Toxin-Antitoxin Modules

**DOI:** 10.1101/146027

**Authors:** Curtis T. Ogle, William H. Mather

## Abstract

Chronic bacterial infections present a serious threat to the health of humans by decreasing life expectancy and quality. Resilience of these populations is closely linked to a small fraction of persister cells that are capable of surviving a wide range of environmental stressors that include starvation, DNA damage, heat shock, and antibiotics. In contrast to inherited resistance, persistence arises from a rare and reversible phenotypic change that protects the cell for one or a few generations. The frequency and character of persistence is controlled in part by the dynamics of numerous toxin-antitoxin (TA) modules, operons with an evolutionarily conserved motif including a toxin that slows cell growth and an antitoxin that can neutralize the toxin. While many such modules have been identified and studied in a wide range of organisms, relatively little consideration of the interactions between multiple TA modules within a single host has been made. Particularly, a multitude of different protein-based antitoxin species are known to be actively degraded by a limited number of shared proteolytic pathways, strongly suggesting interaction via competition between antitoxins for degradation machinery. Here we present a theoretical understanding of the dynamics of multiple TA modules whose activity is coupled through either proteolytic activity, a toxic effect on cell growth rate, or both. We also present a generalizable theoretical mechanism by which a toxic state is tunable by regulation of proteolysis. Such regulation or indirect coordination between multiple TA modules may be at the heart of the flexibility and robustness observed for bacterial persistence.

## I. Introduction

It is well established that many bacteria are capable of collective behaviors which yield survival strategies unavailable to individual cells [1–4]. Bacterial persistence is one such behavior that provides a broad spectrum response to potentially deadly events within the environment. Persistence is characterized by a small fraction of a bacterial population occupying a quasi-dormant (persistent) state in which the cells hardly metabolize or grow. This state provides immediate robustness against a wide range of environmental stressors such as starvation and antibiotic treatment, allowing the small subpopulation to survive events which would kill most normally growing cells [3, 5–11]. With modern medicine, this strategy is indispensable to bacteria that are capable of chronic infection within humans [3, 9, 12]. It has been claimed that as much as 65% to 80% of all bacterial infections are attributable to bacterial persistence [3, 13]. Persistence is also strongly linked to biofilm formation and survival [2, 8, 9], which is both necessary for many healthy processes in humans and the most likely source of hospital borne disease [1].

Toxin-antitoxin (TA) modules have been found to be central in the dynamics of bacterial persistence [3, 7–11, 13–16]. TA modules are small gene networks following a motif that includes two or more genes within an operon, one of which is a relatively stable toxin that slows the growth of the cell when it accumulates, and the other of which is a relatively unstable antitoxin that neutralizes the toxin. While several classes of TA modules have been studied, considerable attention has been given to type II modules, where the antitoxin is a protein that is capable of forming complexes with its associated toxin [7, 10, 11, 15, 17–19]. TA modules are thought to exist in two major states. Normally, TA modules reenforce a state where almost all of the toxin molecules are bound in complex with antitoxin, thus neutralizing the action of toxin. This state is in contrast to a high free toxin state, where a sufficient quantity of free toxin accumulates and results in cell growth arrest characteristic of the persistent state [10, 11]. These two states theoretically are metastable, and this view has led TA modules to be interpreted as bistable switches, where stochastic transitions govern jumping between states [6].

Many bacteria have a surprisingly high number of TA modules (at least 36 within *Escherichia coli* [3, 13, 15] and at least 88 within *Mycobacterium tuberculosis* [2, 3, 7, 13, 16, 17]), though why this is the case has not been well explained [3]. The nonpathogenic organism *Mycobacterium smegmatis* has only two putative TA modules, suggesting that the plurality of TA modules is related to the pathogenicity of some bacterial strains, or that many TA modules play a role in achieving the particularly long periods of dormancy associated with M. tuberculosis [3, 15]. It is thus important for human medicine to understand the interplay of potentially many TA modules in promoting pathogenicity and in particular bacterial persistence. Active proteolytic degradation of the antitoxin protein is equally important for persistence. For *E*. *coli*, it has been shown that removal of either the Lon protease or a number of known TA modules is sufficient to strongly suppress persistence [7, 9]. This suggests that both the presence of specific TA modules and proteolytic degradation are necessary for persister formation. In *E*. *coli*, there are three proteases (Lon, ClpAP, and ClpXP) responsible for more than 70% of ATP-dependent protein degradation within the cell [20, 21]. In particular, Lon is responsible for approximately 50% of defective protein degradation and likely degrades the antitoxins of more than 10 TA modules in *E*. *coli* [7, 9]. There are thus far fewer proteolytic pathways than the number of TA modules, implying that more than one TA module is actively utilizing a given proteolytic pathway.

The effect of processing bottlenecks on biological networks can be understood in the context of queueing theory. Queueing theory is a formalism traditionally used to describe the dynamics of a set of servers that serve a set of customers, but it more broadly offers insight into the effects of bottlenecks on a system [22]. In particular, queueing theory has been used to explain the observation of strong correlations appearing in networks containing molecules competing for enzymatic processing resources [23–26], with these correlations being maximized near the queueing theoretic point of balance in a phenomenon known as correlation resonance [23, 24, 26]. It is conceivable that the positively correlating effect of proteolytic competition (queueing coupling) between the antitoxins of TA modules provides a mechanism by which TA modules might coordinate their effects and shape the persistence behavior. We note as an aside that analogous but distinct queueing effects have previously been proposed to gate persistence by the rare overloading of metabolic bandwidth and subsequent metabolic poisoning [27], though we do not pursue such a mechanism here.

In the following, we support that competition between antitoxins for proteolytic machinery is a route to strong positive coordination of TA modules. While several theoretical investigations have considered the dynamics of a single TA module [10, 11], interactions between multiple TA modules have only more recently been explored [3, 13, 18], and to the best of our knowledge, our work is the first to consider queueing coupling of antitoxins. We examine a custom model with components and interactions motivated by the well-studied TA module *mazEF* [2, 15–17, 28, 29]. Extensive stochastic simulation of this model supports that proteolytic competition robustly leads to strong positive coordination between TA modules, whether in the presence or absence of effects due to cell physiology (growth rate coupling [3, 13]). We verify this phenomenon also in the context of a more robust model for TA modules that leverages a novel return mechanism, which limits the period of toxic activity on the cell. Our return mechanism is based on internally regulated affinity of protease to free toxin, as can be motivated by known regulation mechanisms of protease in response to persister physiology [9, 30], but we emphasize that many alternate return mechanisms are plausible and may indeed depend on the particular organism.

An outline of this article is as follows. Section II presents the details of the quantitative models used. Methods for the simulation and analysis of these models appear in Section III. Results and corresponding discussion are presented in Section IV. Section V provides concluding remarks.

## II. Models

We consider models for one or more identical TA modules coupled using components and interactions that are based on a well-studied TA module found in *E*. *coli* (see Figure 1 for a schematic of this network). Specifically, the stoichiometric ratios of the complexes formed by the toxin and antitoxin and the mechanism of toxicity where the toxin cleaves mRNAs with specific sequences are based on *mazEF* [2, 15, 17, 28, 29]. *mazEF* is of interest for exploring proteolytic coupling, as the antitoxin MazE is degraded by both ClpAP and Lon [2, 15, 18, 28] and could thus indirectly couple additional modules as in Ref. [26]. The toxin MazF inhibits the translation of approximately 90% of all proteins in *E*. *coli* [17] and thus plays an important role in specifying protein production during environmental stress.

**FIG. 1:**
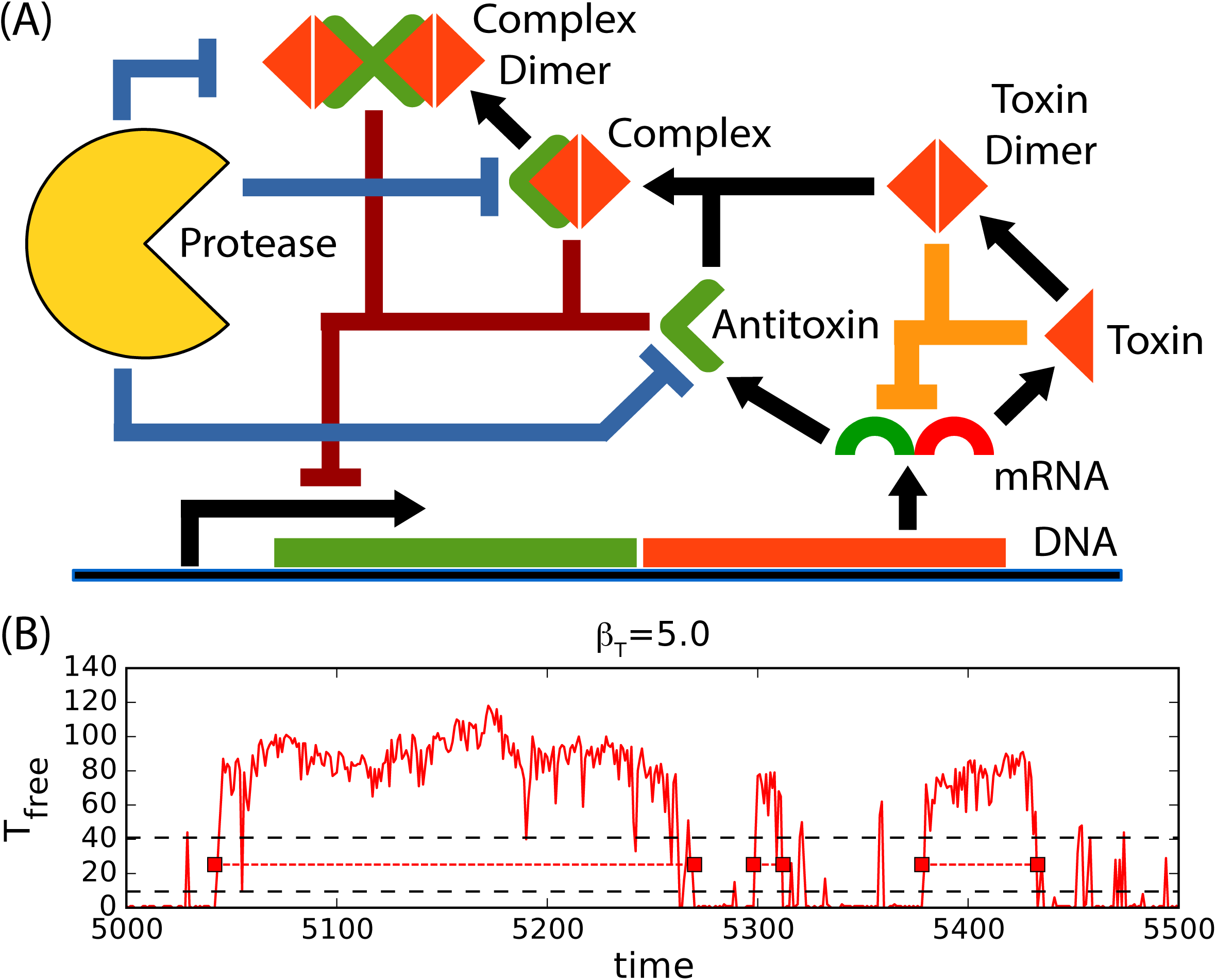
(A) Schematic of our model for a TA module with features similar to the *mazEF* system. A single mRNA is produced encoding both toxin and antitoxin proteins. The toxin protein forms a dimer, which then forms a complex with the antitoxin protein. This complex can also dimerize. The antitoxin, complex, and complex dimer all repress transcription of the mRNA (dark red lines). Proteases can degrade free antitoxin and antitoxin bound in complexes (blue lines). Free toxin and its dimer inhibit translation of both the toxin and antitoxin by cleaving the mRNA encoding them (orange lines). All species are also subject to effective dilution by cell growth and division (not shown). The rate of dilution is dependent on the level of free toxin and its dimer because cell growth is assumed to be slowed by toxin activity. (B) A sample single trajectory for free toxin count *T_free_* = (*T* + *T*_2_) (red lines) of a single isolated TA module, with detected high free toxin events (toxic events) indicated (dashed red lines and solid boxes). Dashed black lines represent high and low thresholds used to detect events (see Methods Section IIIB). The trajectory and its thresholds are derived from a simulation of 10 time units. Parameters are *β_T_* = 5.0 and standard otherwise (see Models Section II).

A general model for *M* such identical TA modules interacting via *N* identical proteolytic pathways and via toxic effect on a shared cell growth rate is presented in Section II A. In Section IIB we describe several special cases of the general model for which results are provided. We provide a modified version of this model to include a return mechanism in Section IIC. A description of the algorithm and software used is provided in Section III A. The algorithm for the detection of high free toxin states (persister-like states termed events) is described in section IIIB, and measurement of event statistics is described in section IIIC.

### A. Model of Coupled TA Modules

We primarily use models that are special cases of a more general model for *M* identical TA modules interacting via *N* identical proteolytic pathways (inspired by the model in Ref. [26]) and via toxic effects on the cell growth rate. Each module, based on the well-studied *mazEF* module as found in *E. coli* (see Figure 1), tracks the counts for toxin-encoding and antitoxin-encoding mRNA’s (*t_j_* and *a_j_*, respectively), the toxin and antitoxin proteins (*T_j_* and *A_j_*, respectively), the dimer of the toxin protein (*T*_*j*2_), the complex formed by the toxin dimer and antitoxin (*C_j_*), and the dimer of the complex (*C*_*j*2_). The subscript *j* ranges from 1 ≤ *j* ≤ *M* and denotes the TA module with which these components associate. Complex formation results in a stoichiometric ratio of two toxin proteins to one antitoxin protein, as observed in the *mazEF* module [15, 28]. Both toxin and antitoxin mRNAs are cleaved by free toxin and its dimer. The degradation rates of free antitoxin, antitoxin bound in complexes, and free toxin are represented using functional forms similar to those used in [23, 26]. All species of the *j*^*th*^ module are diluted at common rate *Γ_j_*, representing the process of cell growth and division. The functional dependence of *Γ_j_* on the system state encodes the effect of free toxin on growth rate. Though we in principle allow each *Γ_j_* to be independent of other TA modules, realistic models will have a common value for each *Γ_j_* to reflect the growth rate for the host cell.

Our base model is implemented by several stochastic reactions, which are designed to allow for the model to have molecular level detail, and which are intended to provide a generic platform for understanding TA modules rather than providing a particular fitted model. This model also does not yet include our implementation of a return mechanism (discussed in more detail in Section IIC). The reactions for the model are as follows, where we assume mass action kinetics when defining the propensities of reactions [31]. Transcription of a shared mRNA for toxin and antitoxin is modeled as the reaction

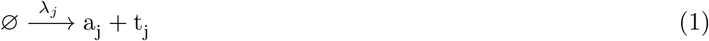

where the function

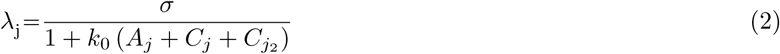

encodes how complex and free antitoxin represses transcription, with *σ* the maximal transcription rate. Note that we model *a_j_* and *t_j_* as independent entities, which allows for them to be degraded at different rates, as might be expected for prokaryotic degradation [32]. Also, we do not include crosstalk due to transcriptional or translational coupling between modules. Degradation of mRNA molecules follows from the reactions

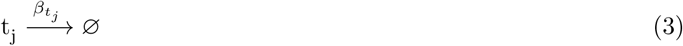

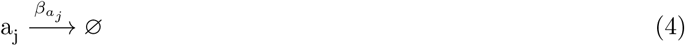

where the functions

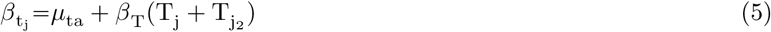

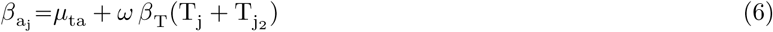

encode both basal degradation rate *µ_ta_* and accelerated degradation due to toxin. The parameter *ω* allows for differential degradation of antitoxin and toxin transcripts by the toxin T_j_, e.g. due to a different number of cut sites in the transcript. Translation of proteins follows from the reactions

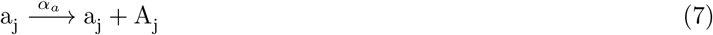

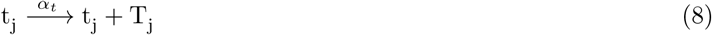

where *α_t_* is a constant. We relate translation rates by

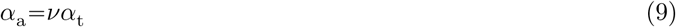

where *v* scales the relative translation rate of antitoxin to that of the toxin (expected to be genetically encoded by the transcript). This parameter is one of the few very sensitive parameters we identified in our model, and it is set by the need for cells to consistently produce enough antitoxin protein to deactivate toxin protein [2]. Formation of toxin dimers and complexes follows from the reactions

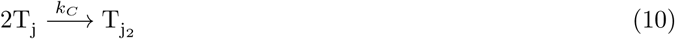

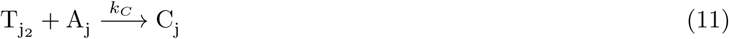

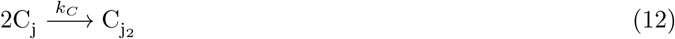

for which we assume irreversible (tight) binding, for simplicity. It is worth noting that we use the convention that the reaction 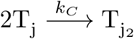 occurs with velocity (1/2) *k_C_ T_j_* (*T_j_* – 1), and the reaction 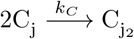 occurs with velocity (1/2) *k_C_ C_j_*(*C_j_*–1). Proteolytic degradation of antitoxin and other species by protease is modeled by multiple reactions

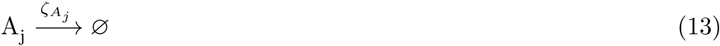

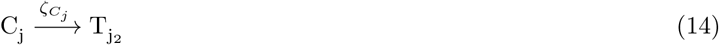

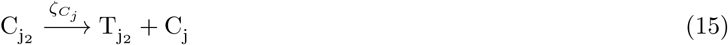

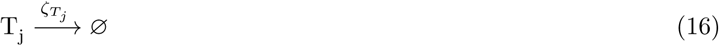

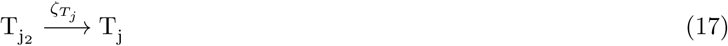

Where

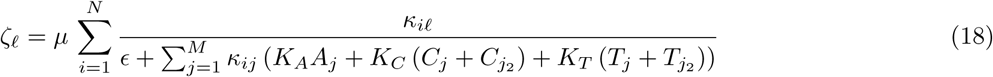

and

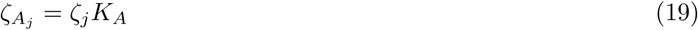

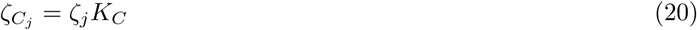

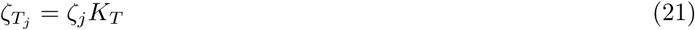

define reaction rates for competitive proteolytic degradation. The coefficients *κ_ij_* will define with what affinity multiple proteases target multiple TA modules [26], while the coefficients *K_a_, K_c_*, and *K_T_* define with what affinity antitoxin, complex, and toxin are degraded, respectively. The value of e sets the overall affinity of substrate to protease. Note that we use the convention that the reaction velocities for Eqs. 13–17 (as in all our reactions) are determined by multiplication of their reaction rate functions by their respective mass action terms. Finally, all species are removed from the system by dilution (cell growth and division), which leads to a reaction for all species (generically labeled *Z_j_*) in a TA module

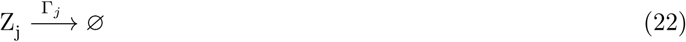

with a dilution rate

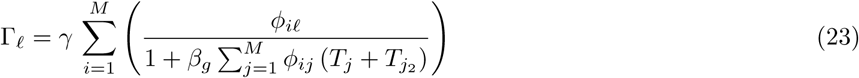

that is sensitive to free toxin levels. The coefficients *φ_ij_* allow for the following scenarios

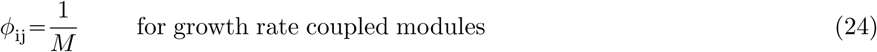

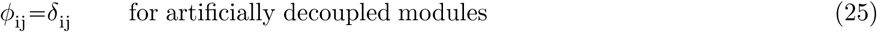

with *δ_ij_* the Kronecker delta (1 if indices are equal and 0 otherwise).

Values for the above parameters are as follows unless otherwise specified *σ* = 20.0, *k_0_* = 0.05, *v* = 1.6, *α_t_* = 200.0, *k_C_* = 1000.0, *µ_ta_* = 5.0, *ω* = 0.2, *β_T_*= 20.0, *µ* = 100.0, *K_a_* = 1.0, *K_C_* = 0.1, *K_T_* = 0.0, *∈* = 0.01, *γ* = 1.0, *β_g_=* 0.25. This parameter set leads to infrequent high toxin (persister-like) events for a single TA module. This frequency is usually much higher than that seen experimentally (approximate persister frequency range 10^−6^ to 10^−4^ in experiments), but we have chosen a high persister frequency to efficiently explore the dynamics of our model using direct stochastic simulation only. Extension of our methods to much lower persister frequencies is planned for future work.

Parameter values in our model were motivated by experimental observations and theoretical flexibility when possible. Our choice of *ω* = 0.2 follows from a simple analysis that compares the number of toxin cut sites (ACA) on the toxin vs. antitoxin segments of the mRNA for the *mazEF* TA module. Our choice of *v* > 1 follows from the general observation that antitoxin protein should be produced at a higher rate than toxin to ensure proper function of the TA module [2]. Our choice of small e ensures generally high antitoxin affinity to protease [26], which is likely the case if antitoxin is degraded quickly and efficiently in the cell. Our choice of *γ* = 1.0 for the maximum growth rate of cells was achieved by rescaling time (this can always be done). Parameter values were also motivated by a decision to investigate a model with specific properties. Our choice of large *k_C_* encodes the assumption that toxin monomers rapidly dimerize, that free toxin dimers rapidly bind to antitoxin to form complexes, and that toxin-antitoxin complexes can rapidly dimerize. Our choice of *K_A_* = 1.0, *K_C_* = 0.1, *K_T_* = 0.0 reflects that free antitoxin is more likely to be degraded than antitoxin in complexes (this is apparently not critical, as will be supported in Fig. 4), and that toxin is relatively unlikely to undergo proteolytic degradation. The overall scale for parameter values associated with transcription and translation was adjusted to ensure that production and degradation reactions were sufficiently rapid relative to dilution. Remaining parameters were adjusted self consistently to achieve a system with sufficiently high persister frequencies that are readily accessible by direct stochastic simulation.

A number of other potential modes of coupling were not included in our model for simplicity. These include the effect of toxin activity directly on transcription rates due to altered metabolic state of the cell, targeting of a TA module’s mRNA by another TA module’s toxin, and transcriptional crosstalk between TA modules at the promoter level.

### B. Particular Models without a Return Mechanism

The majority of this work uses the above model, which exhibits bistable behavior for a variety of parameter values. We begin our exploration with the simplest variation of the general model, where *M = N* = 1. Figure 1 contains a basic schematic of this situation. No proteolytic or growth rate coupling is investigated here. Rather, the aim is to establish a baseline understanding of the *mazEF-like* module. Results of this model are discussed in Section IV A.

We next consider the case where *M = N* = 2. Figure 2 contains a basic schematic of this situation. This model permits exploration of the effects of both coupling via proteolytic competition and via toxic effect on a shared cell growth rate. To this end, we impose a parameterization of the proteolytic affinities of each module, *κ_ij_*, as in Ref. [26].

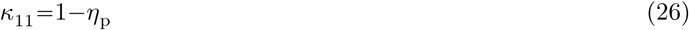

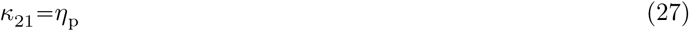

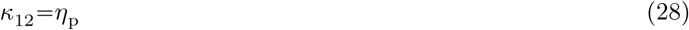

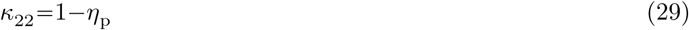

The strength of proteolytic coupling is encoded in the parameter *η_p_*. This additional parameter is bounded below by 0, where coupling is nonexistent, and above by 0.5 for maximal proteolytic coupling. As *η_p_* is increased above 0.5, the identities of the associated proteolytic pathways are effectively exchanged due to symmetry in the representations of *ζ_j_*. Results of this model are discussed in Section IV B.

**FIG. 2:**
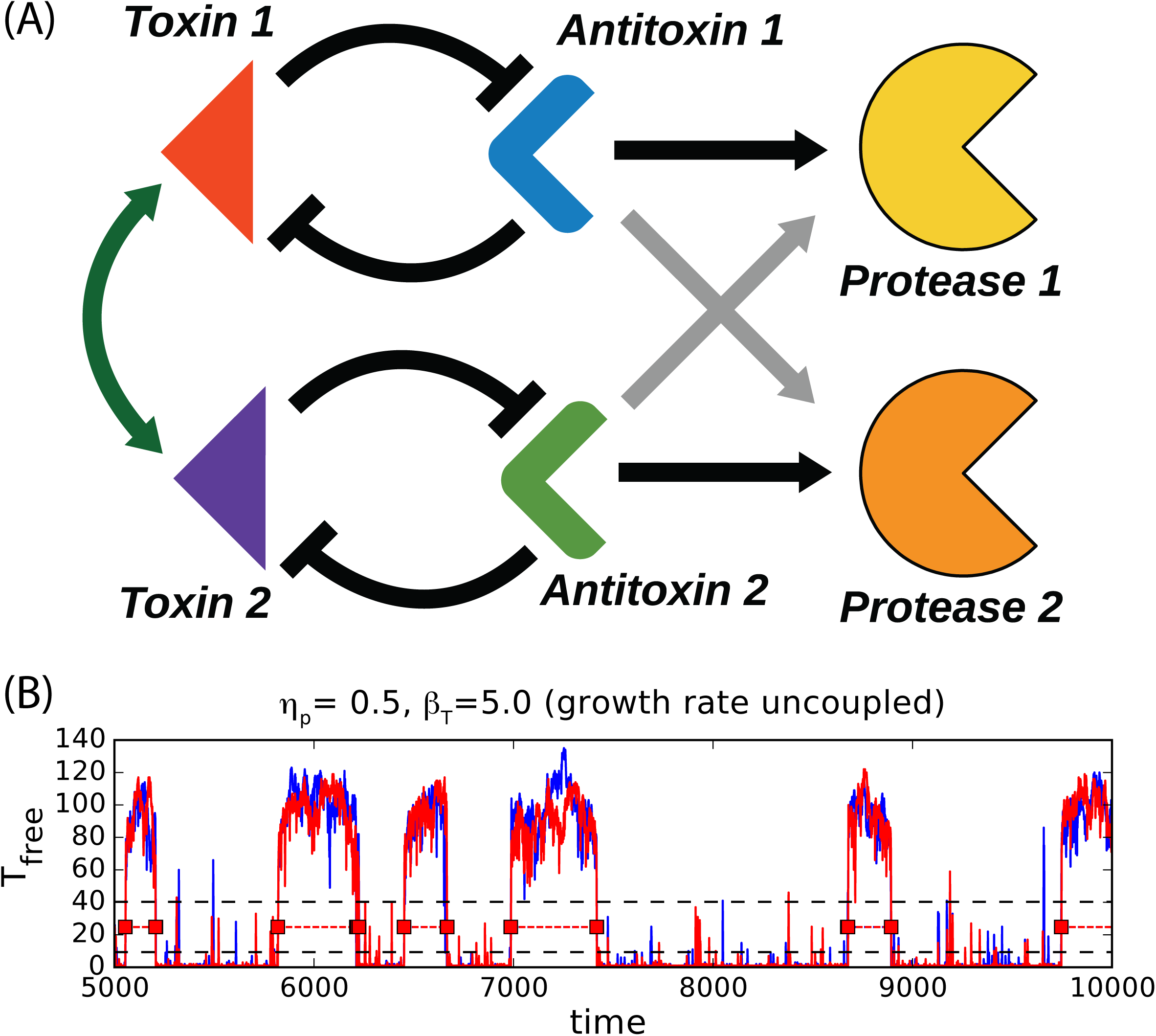
(A) With the addition of an identical *mazEF*-like module and a proteolytic pathway, the effect of proteolytic coupling can be measured. There are now four distint proteolytic actions, where either pathway may process the antitoxin of either module. In the uncoupled case, the pathways shown by grey arrows would not be present. Maximal coupling requires that antitoxins do not prefer either proteolytic pathway over the other (both pathways effectively constitute a single pathway) [26]. The antitoxins are shown to repress their associated toxins because of the neutralization concomitant with complex formation (and autorepression of the operon). The toxin is shown to repress the antitoxin as it inhibits translation preventing the production of additional antitoxin (and toxin). With proteolytic coupling, the toxins of each module are coupled transitively (green arrows) because of antitoxin coupling via proteolysis. Although artificial, the effective separation of dilutive pathways presents a similar picture, except that toxin-toxin interaction is more direct as both toxins are diluted, as opposed to the strictly transitive coupling of toxins owed to proteolytic competition. (B) A representative trajectory for TA modules with perfect proteolytic coupling (*η_p_* = 0.5) but no growth rate coupling or other coupling. It is evident that events for the two modules are tightly coordinated by this coupling. Parameters are *η_p_* = 0.5, *β_T_* = 5.0, no growth coupling (Eq. 25), and standard otherwise (see Models Section II).

### C. Particular Model with a Return Mechanism

As discussed in Section IV C, reliance on a bistable model for persistence leads to a number of negative consequences, including a potentially low rate to return to a normal (low free toxin) state. A system that is instead excitable (noise-induced transition to the toxic state, but more reliable transition to the normal state) can be created by regulating the mean lifetime of the toxic state. There are a multitude of potential return mechanisms that could lead to this behavior, but we investigate for definiteness only one such mechanism: a molecule is produced during the toxic state that increases the affinity of free toxin (both monomeric and dimeric) towards proteolytic degradation. This mechanism is motivated by known regulation of proteolytic pathways affecting persistence [9, 30]. For brevity, We will consider this mechanism only in the case of two coupled TA modules that include perfect growth rate coupling (*φ_ij_* = 1/2 in Eq. 24), but effectiveness of the return mechanism holds for other parameter regimes.

For this, we introduce an additional molecule (e.g. protein) *X* with precursor (e.g. mRNA) *x*. The identities of these molecules are not particularly important, but it is important that *X* is created or at least active when the system has entered a toxic state. The molecule *x*, which is only produced in the presence of free toxin (both monomeric and dimeric), is produced at rate *σ_x_*, while *X* is produced by *x* at rate *α_X_*. This is implemented by the reactions

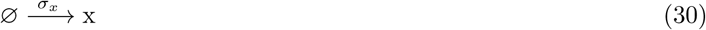

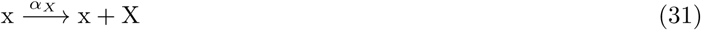

where the production rate of *x* is state dependent

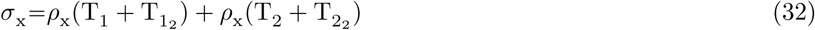

Both *x* and *X* are diluted at rate *Γ_1_* for simplicity via the reactions

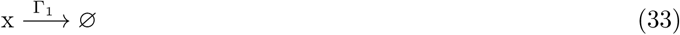

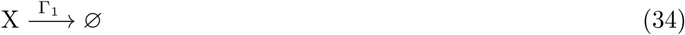

where recall that *Γ*_1_ and *Γ*_2_ are identical. We suppose the protein *X* permits the degradation of toxin by increasing *K_T_*, which in turn allows toxin to be degraded by protease. The strength this effect is encoded by *β_X_* by substituting

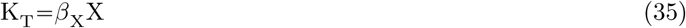

Thus, in the absence of *X* or if *β_X_* = 0, free toxin is not actively degraded by protease. Typical values for the return mechanism are *ρ_x_* = 0.05, *α_X_* = 0.01, and *β_X_* = 10^−7^ which lead to notably reduced mean event lifetimes.

## III. Methods

### A. Simulation Algorithm

All simulations were performed using a custom implementation of the Gillespie algorithm [31] in Python leveraging optimizations made possible by the Cython library [33]. Libraries from the Scipy stack [34] were used for analysis. These simulations were primarily executed and analyzed using a custom Python package, available at www.github.com/ctogle/modular.

### B. Event Detection

To analyze the likelihood that one or more TA modules are in a sufficiently high concentration (toxic) or sufficiently low concentration (normal) free toxin state, we automatically identify windows of time, labeled events, with high free toxin (monomeric and dimeric). The following details a particular algorithm for event detection that produced reasonable results in our study.

Event detection for the *j*^*th*^ module only considers the count of total free toxin *T_j,free_* = (*T_j_* + *T*_*j*2_) after an initial transient (ignoring the first 0.1% of each long realization). To scale our definition of a high free toxin state, we define 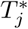 as the 99.9^th^ percentile of *T_j,free_*. We initially defined 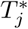 as the maximum value of *T_j,free_* over time, but this definition was avoided because the maximum is not a stable statistic (the expectation value of the maximum continues to increase for longer duration trajectories). The algorithm uses 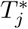 to define a low transition threshold (*T_low_*) and the high transition threshold (*T_high_*) by the following definition

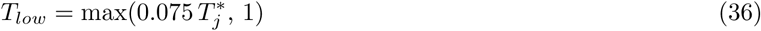

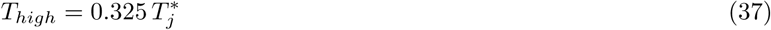

The difference of these thresholds must be greater than or equal to 4, i.e. *T_high_* – *T_low_* ≥ 4, or else no events are detected for a trajectory. The numbers 0.075 and 0.325 are somewhat arbitrary and may be adjusted in other studies, but we keep these numbers fixed in this study. Subsequent studies may consider a more general algorithm based on clustering analysis to identify these thresholds.

Identification of events proceeds in two phases: identification of primary events and filtering of events. Primary events are windows of time defined by consecutive time points all satisfying *T_j,free_* ≥ *T_low_*, with at least one such time point satisfying *T_j,free_* ≥ *T_high_*, and with the requirement that this window of time has a boundary consisting of two time points satisfying *T_j,free_* < *T_low_*. The collection of primary events already represents a useful tool for the statistical analysis of our model, but we opted to filter these events to remove short isolated events and to merge events separated by short gaps of time. Our detailed algorithm for this filtering is controlled by a single filtering parameter *n_f_*, which we fix *n_f_* = 5 time units (note that simulations are sampled every 1 time unit). Filtering is as follows

1. Consider an ordered list of events *L_E_* that will be the final list of events after filtering. When it exists, the last event in this list is labeled *E_l_*.
2. *L_E_* initially contains only the earliest primary event with more than *n_f_* data points. Define *E_c_* as the immediately following primary event. If *L_E_* is empty or *E_c_* does not exist, abort filtering.
3. Determine if the data points in the gap between *E_l_* and *E_c_* satisfy either of the following: (a) the gap contains fewer than n_f_ data points, or (b) mean free toxin in the gap exceeds *T_high_* (this latter condition is very rare in our simulations). If so, modify the event *E_l_* to be the merger of events *E_l_* and *E_c_* (the new event is the the window of time defined by the start of *E_l_* and the end of *E_c_*). If not, append the event *E_c_* to the list *L_E_* if it contains more than *n_f_* data points.
4. If possible, update the event *E_c_* to be the event immediately following the current event *E_c_*, then repeat Step 3. Otherwise, continue to Step 5.
5. Remove events in the list *L_E_* that include points within *n_f_* data points of the first and last time points of the data.

The net result of this filter is a list *L_E_* of events used for analysis.

In reporting results, we exclude simulations with fewer than 100 filtered events for any TA module. This is to avoid statistically unreliable data.

### C. Event Measurements

The dynamics of single TA modules are characterized by three primary statistics: number of events, event probability, and event duration. Event probability *P_i_* for the *i*^*th*^ TA module is found by measuring the fraction of time points in a trajectory belonging to events of the *i*^*th*^ TA module. Mean event duration for the *i*^*th*^ TA module is the total time spent in events of the *i*^*th*^ TA module divided by the number of measured events for this TA module.

Coupled TA modules can produce tightly coordinate events. There are a number of approaches to measuring this behavior, and we chose a conditional probability measure. Conditional event probability *P_i_*_|_*_j_* is defined by *P_ij_*/*P_j_*, where the joint event probability *P_ij_* for TA modules i and j is the fraction of time points simultaneously belonging to an event in both TA modules. If TA modules i and j are statistically independent, then *P_ij_* = *P_i_P_j_*, which implies *P_i_*_|_*_j_* = *P_i_*. However, *P_ij_* can approach a value of 1 for TA modules with highly correlated behavior.

## IV. RESULTS AND DISCUSSION

Our investigation reveals that our model for a single TA module without a return mechanism operates essentially as a bistable system, largely owing to the relative instability of antitoxin protein to toxin protein, where we loosely use the term “toxin” here to include both toxin monomers and dimers. Noise allows our model to spontaneously switch between two metastable states, a low free toxin state (representing a cell in a normal metabolic state) and a high free toxin state (representing a cell in a persister-like state). Our choice to include enzymatic degradation of proteins allows for coupling of multiple TA modules through competition (queueing coupling) of antitoxins for proteases, the effect of which is a rapid positive coordination of the TA modules, such that the presence of an event (period of high free toxin) for one TA module strongly predicts a simultaneous event for another TA module. This effect robustly occurs for a wide range of proteolytic coupling. Inclusion of a return mechanism in our model changes the model’s qualitative nature so as to be better described by an excitable system, where events have a short though still stochastic lifetime relative to the mean time between events. Queueing coupling continues to lead to strong positive coordination in the presence of this return mechanism. These observations are derived from the analysis of direct numerical simulations of particular models detailed in this article, but our findings are consistent with a range of analytic, numerical, and experimental results showing the positively correlating effects of queueing coupling [23, 24, 26, 35, 36].

### A. Single TA Module Model

We first investigated the behavior of our model for an isolated TA module, with the goal of understanding how proteolytic queueing influences the dynamics of multiple coupled TA modules. The temporal behavior of this model for the parameters considered leads to essentially bistable behavior, where the system transitions between metastable states either with low levels of free toxin protein or with high levels of free toxin protein (see Fig. 1). We identify toxic state events by considering sufficiently long windows of time in the high free toxin state (see Section IIIB). Correlations between the counts of components and events (see Fig. 3) suggest that events correspond to periods of high free toxin counts, as expected, but also correspond to reduced counts for all of antitoxin-toxin complex, mRNA for antitoxin and toxin, and free antitoxin.

**FIG. 3:**
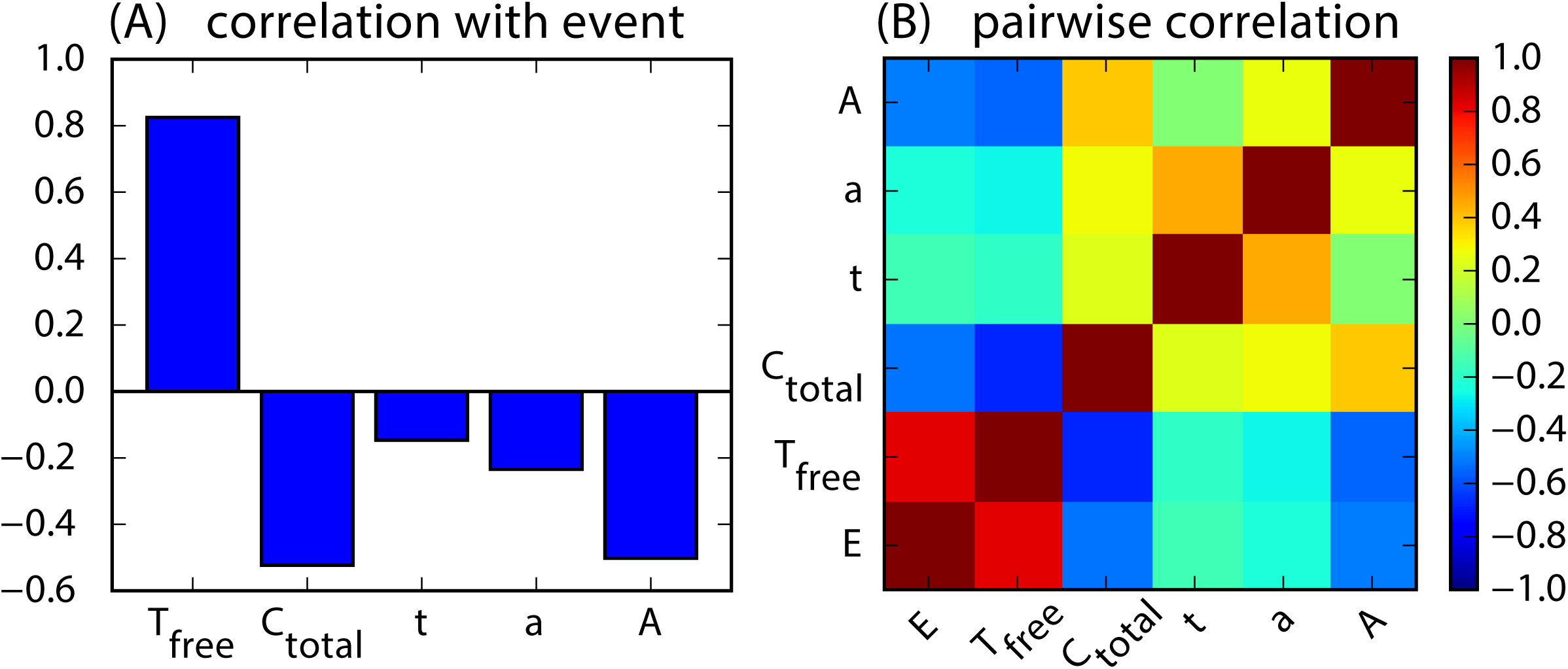
(A) Representation of the higher dimensional switching dynamics of a single isolated TA module without a return mechanism that is simulated for 10^6^ time units. Shown is the Pearson correlation coefficient between events and the counts of free toxin (*T_free_* = *T* +*T*_2_), total complex (*C_total_* = *C* + *C*_2_), mRNA (*t* and *a*), and free antitoxin (A). Correlation coefficient is calculated by first defining a time-dependent quantity *E* that is 1 during an event and 0 otherwise, then by taking the standard correlation coefficient between *E* and other quantities. (B) Similarly for the pairwise correlation coefficient between all these quantities. Parameters are standard (see Models, Section II).

Metastability of these two approximate states was explored by scanning parameters. In the low free toxin state (analogous to the normal metabolic state of cells), metastability of the state is expected to be ensured by the maintenance of sufficiently high antitoxin count relative to toxin count, thus ensuring the toxic effect of toxin is neutralized [10, 11]. The 2:1 ratio of toxin monomer to antitoxin binding in our model predicts that a relative translation ratio *ν* > 0.5 would produce sufficient antitoxin. However, since antitoxin is expected to be degraded at a higher rate than toxin, the value of *ν* required for metastability of the low free toxin state is likely required to be larger than the approximate bound of 0.5. The stabilizing effect of sufficiently large *ν* was demonstrated by significantly decreasing event probability (increasing probability to be in the low free toxin state) with increasing *ν* (see Fig. 4A). Increasing *ν* also decreased mean event time (decreased time in the high free toxin state) (see Fig. 4B). For our study, we chose *ν* = 1.6 as a reasonable value for this parameter.

**FIG. 4:**
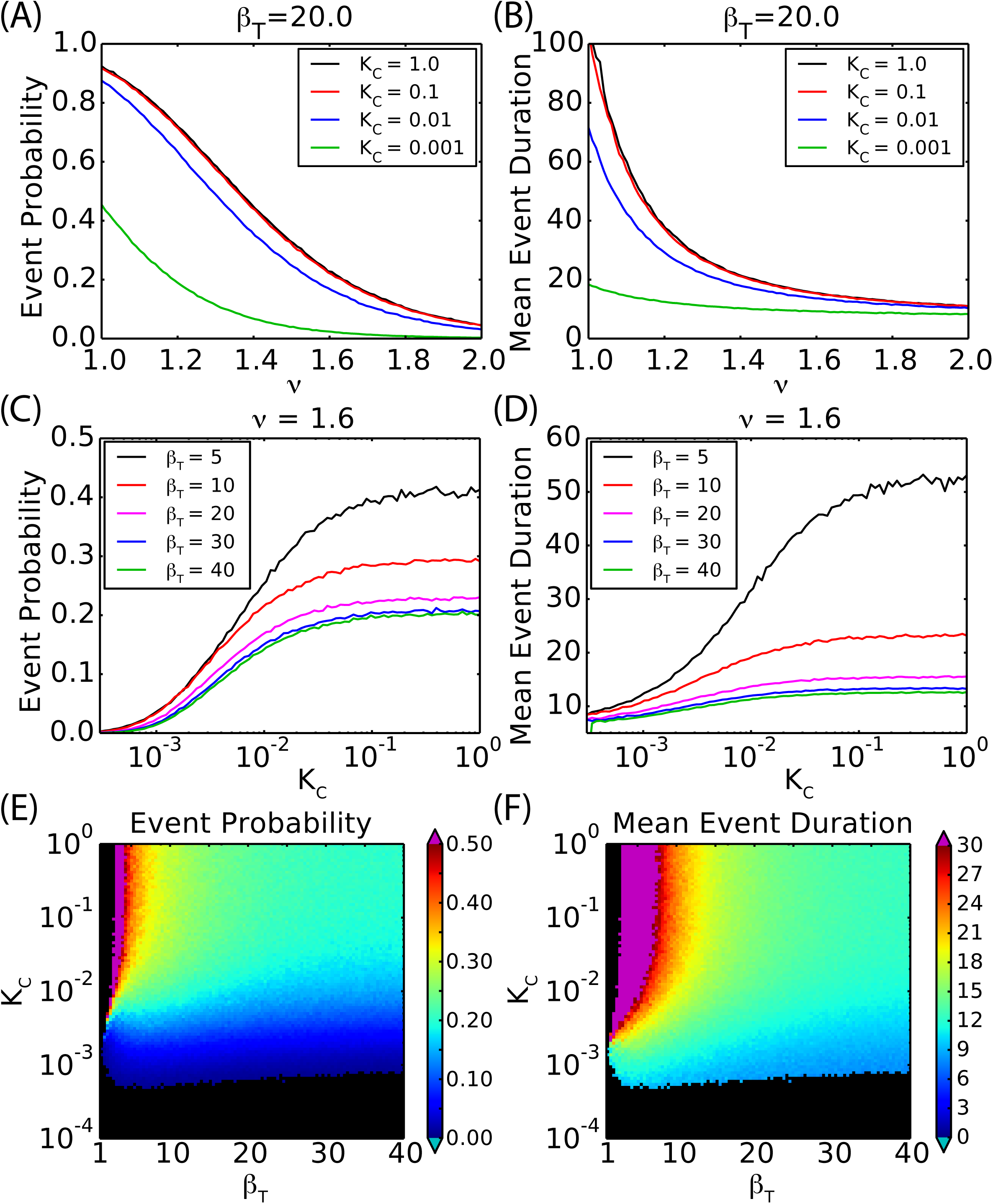
Statistics for a single isolated TA module without a return mechanism for scans of parameters *v, K_C_*, and *β_T_* (see Fig. 1B for a single trajectory). Statistics in (A) through (D) show results using a trajectory of 10 time units for each data point. In these scans, it is seen that decreasing *v*, increasing *K_C_*, and decreasing *β_T_* are associated with increasing event probability and mean event duration. Statistics in (E) and (F) show results scanned more densely over two parameters using trajectories of 10 time units, with the consequence that shorter trajectories lead to fewer events. Pixels shaded black were determined not measurable, primarily due to low event frequency (100 events minimum required, see Event Detection, Section IIIB). Parameters are specified in the figure and standard otherwise (see Models, Section II).

We hypothesized that the system might also be sensitive to the relative magnitude of degradation affinity *K_c_* for complex-bound antitoxin as compared to the affinity *K_A_* of free antitoxin. Our results support that that if *K_A_* = 1.0 and *∈* = 10^−2^ (see Eq. 18), then system statistics were relatively stable if *K_c_* > ∈ (see Fig. 4A–D), which is consistent with a parameter regime where absolute substrate affinity to protease is high [26]. The choice of *K_c_* ≪ *∈* instead led to increased chance to be in the low free toxin state, as expected. We chose to focus on the high affinity parameter regime by choosing *K_c_* = 0.1 as a typical value for this parameter.

Fluctuations can eventually destabilize the low free toxin state inducing transition into the high free toxin state, where toxin is largely unbound to antitoxin and is permitted to actively degrade mRNA transcripts and slow cell growth and division. Toxic activity thus leads to at least two positive feedbacks: degradation of antitoxin mRNA and reduction of toxin dilution rate, the former of which reduces expression of a repressor of toxin activity, and the latter of which allows for buildup of free toxin in the cell. These positive feedbacks enforce maintenance of the high free toxin state. However, toxin activity also asserts a negative feedback: degradation of toxin mRNA. A key parameter that influences the strength of these feedbacks is toxin RNase activity, set by the parameter *β_T_*. Indeed, we found that scanning values for *β_T_* strongly influenced event statistics (see Fig. 4). In particular, very low values for *β_T_* led to the system becoming “stuck” in the high free toxin state, since only positive feedback due to growth rate reduction remains from the above three feedback mechanisms. Larger values of *β_T_* reliably led to regular events.

Our above investigation led us to explore parameter values near our standard parameter values (see Models, Section II) when investigating multiple couple TA modules.

### B. Double TA Module Model

Having established that our model for isolated TA modules exhibits stochastic switching between two states that roughly correspond to normal and toxic states, we then considered two identical but distinguishable TA modules that are coupled (see Fig. 2). We predict that relatively modest coupling between these modules could dramatically influence coordination between their events. Intuitively, transitions between metastable states in bistable systems are thought to be exponentially sensitive to parameters in general based on the formalism of transition state theory and similar large deviation theories [37, 38]. Coupling between modules could then lead to strong positive correlation (an event for one TA module promotes an event for the other TA module) or even strong negative correlation (an event for one TA module greatly diminishes the chance of an event in another module).

This picture for probabilistic TA module coupling led us to measure the probability *P*_2|1_ of an event for one TA module (labeled TA module 2) conditional on an event for another TA module (labeled TA module 1). By symmetry of our system, statistics are identical if we exchange the labels 1 and 2. A value of *P*_2|1_ approaching 1 ensures that one event is highly likely given another event. More precisely, if *P*_2|1_ > *P*_2_, where *P*_2_ is the event probability for TA module 2, then this corresponds to events that are positively coordinated, i.e. events for TA module 1 predict events for TA module 2. If *P*_2|1_ < *P*_2_, then events are negatively coordinated, i.e. an event for TA module 1 instead leads to a decreased chance for TA module 2 events. It is of note that if the baseline event probability *P*_2_ approaches 1, as might be the case for a system that has become stabilized towards the high free toxin state (long events separated by short intervals between events), then *P*_2|1_ approaching 1 by itself does not necessarily imply strong positive coordination between events.

Two modes of coupling were considered when measuring the probabilistic coordination between TA modules. The first was proteolytic coupling, or queueing coupling, where antitoxins compete for common proteases. Antitoxin affinity for the less preferred protease is modulated by the parameter *η_p_*, such that proteolytic coupling is absent when *η_p_* = 0.0 and strongest when *η_p_* = 0.5. Our prediction is that when one TA module is in a high free toxin state, then the free bandwidth of the preferred proteolytic pathway taken by this antitoxin has been increased, which then allows for more rapid degradation of antitoxins (free and in complex) associated with other TA modules, which in turn promotes switching of these other TA modules to a high free toxin state. Hence, we predict proteolytic coupling leads to positive event coordination, i.e. an event for one TA module promotes an event for other TA modules. This positively correlating effect would be similar to the phenomenon of correlation resonance studied previously [23]. The second mode of coupling considered was growth rate coupling, where the dilution rate of all cellular molecules is reduced by the free toxin’s effect on cell growth and division. We predict growth rate coupling leads to positive coordination similar to proteolytic coupling, since low dilution rate is apparently an important stabilization factor for the high free toxin state. Though growth rate coupling can in principle be continuously parameterized, we only considered the cases where growth rate coupling was either completely absent (Eq. 25) or completely present (Eq. 24).

Measurement of system behavior for coupled TA modules revealed that either proteolytic coupling or growth rate coupling could lead to substantial positive coordination between TA modules, though proteolytic coupling led to stronger coordination in our model. In the absence of growth rate coupling (see Fig. 5A,B,C), a lack of proteolytic coupling *(η_p_* = 0.0) sensibly produced a conditional probability equal to the event probability, as would arise from statistically independent TA modules. Increasing *η_p_* to 0.5 led to a lower event probability and a conditional probability much higher than the event probability, indicating a strong positive coordination between events of different TA modules. Strong coordination persisted roughly for the range *η_p_* > *∈*, where *∈* = 10^−2^ scales the absolute affinity of substrate to protease. This result is consistent with earlier work showing that absolute affinity sets an important boundary for system correlations [26]. Models that included growth rate coupling (see Fig. 5D,E,F) exhibited similar results but with stronger coordination between events. These results support that even in the presence of growth rate coupling, proteolytic coupling can lead to substantial additional coordination between events.

**FIG. 5:**
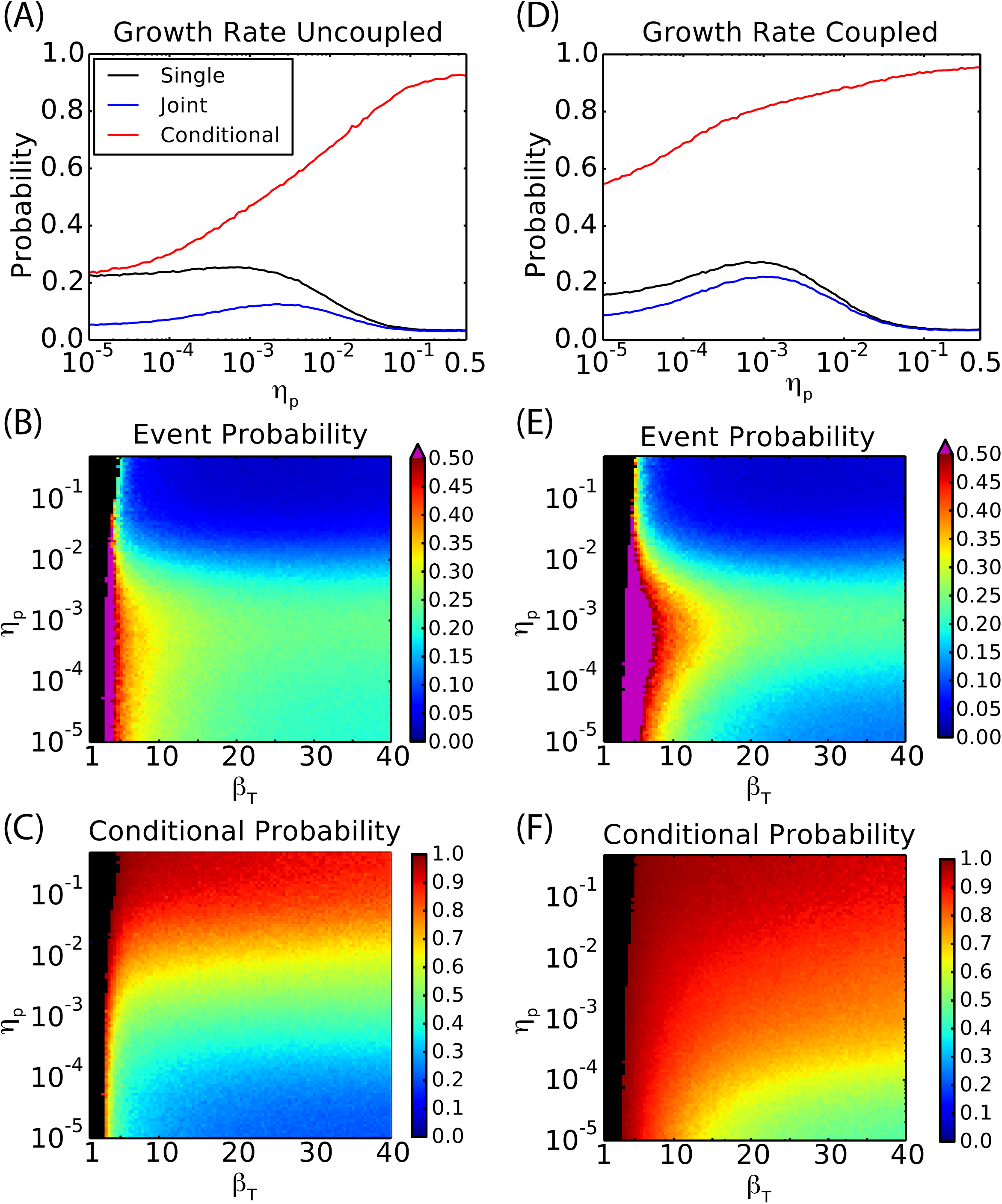
Coordination between TA modules was studied in the case where two TA modules couple to one another by either proteolytic or growth rate coupling (see Fig. 2B for single trajectories showing correlation). (A) For trajectories of duration 10 time units for a system without growth rate coupling (Eq. 25) and with *β_T_* = 20.0, the event probability for a single TA module 1 (black line), the joint event probability for TA modules 1 and 2, and the event probability of TA module 1 conditional on an event for TA module 2 were computed as a function of the proteolytic coupling parameter *η_p_* (see Section IIIC for definitions of these probabilities). It was seen that a wide range of proteolytic coupling strengths led to strong coordination between TA modules, as indicated by their mutual conditional probabilities approaching 1. (B) and (C) show event probability and conditional probability, respectively, for shorter trajectories of length 10 as the parameters *η_p_* and *β_T_* are scanned. (D), (E), and (F) are the same as (A), (B), and (C), respectively, but for a system with perfect growth rate coupling (Eq. 24). Additional growth rate coupling was seen to overall increase the conditional probability while lowering the single event probability, supporting enhanced coordination between TA modules. Parameters are specified in the figure and standard otherwise (see Models, Section II).

### C. Single TA Module Return Mechanism Models

Robustness of coordination due to proteolytic coupling was tested for a modification of our model to include a return mechanism that limits the lifetime of a high free toxin state by increasing the affinity of toxin to proteolytic degradation in response to the accumulation of free toxin (see Section IIC for details). This negative feedback on free toxin is only one of many possible return mechanisms, but existence of a return mechanism ensures cells can reliably reset to a normal physiological state in a reasonable time (see Fig. 6A,B). Addition of a return mechanism to the original bistable model for TA modules leads instead to an excitable system, where transition to the high free toxin state can be slow and highly stochastic, while transition away from the high free toxin state can be faster and often more regular. In principle, addition of a return mechanism can even lead to a model that is oscillatory (results not shown), but such behavior does not seem to be relevant for the typical behavior of TA modules.

**FIG. 6:**
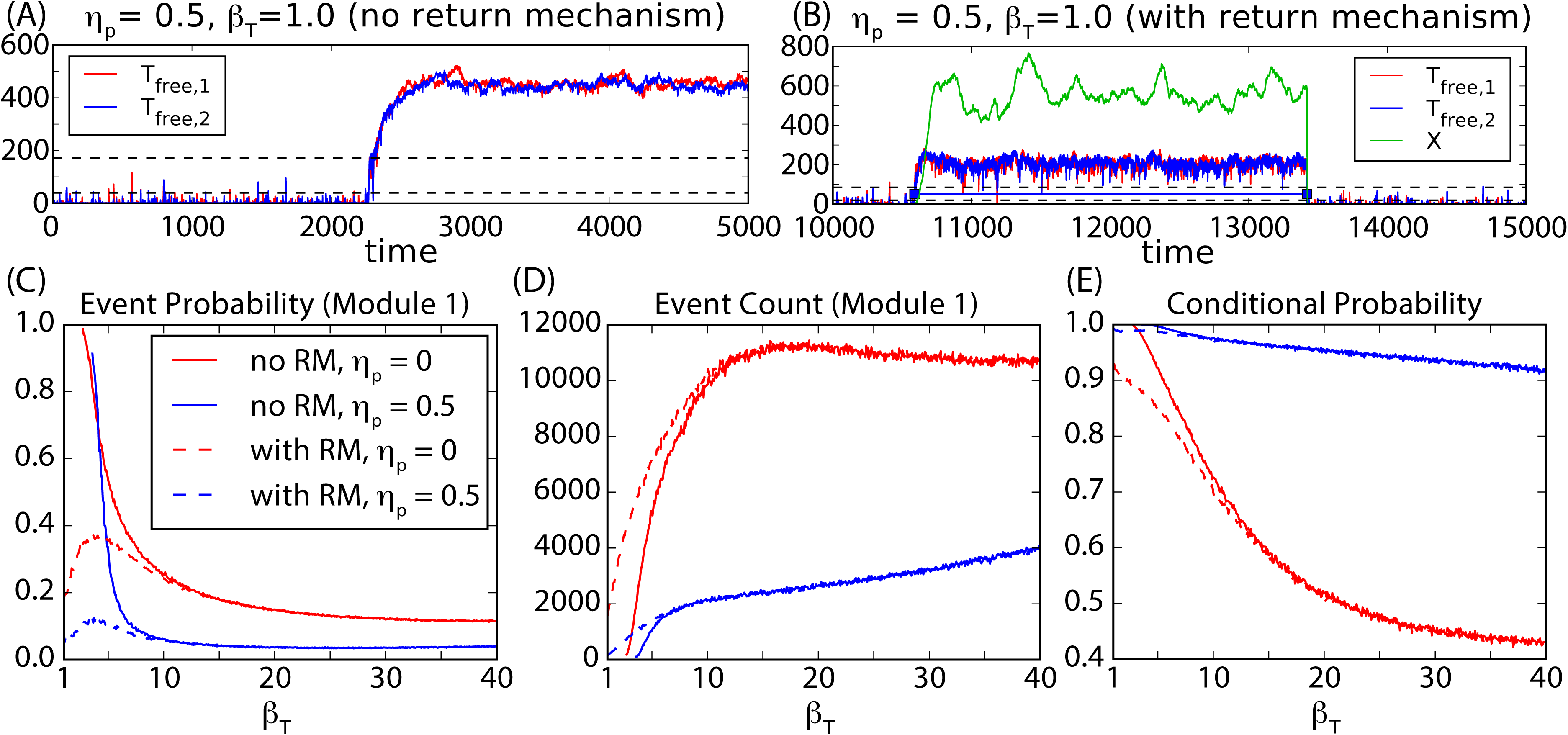
Inclusion of a return mechanism for a model with growth rate coupling leads to increased robustness while maintaining our results for positive coordination. (A) For a model with growth rate coupling (Eq. 24) and without a return mechanism, the choice of parameters *η_p_* = 0.5 and *β_T_* = 1.0 leads to simulations that switch to a high free toxin state but fail to return to a low free toxin state by the end of a 10 time unit simulation. (B) The same model with return mechanism added leads to a series of finite duration events. (C), (D), and (E) show event probability, event count, and conditional probability, respectively, for this model as a function of *β_T_* and for different values of proteolytic coupling strength *η_p_*. Statistics for models without a return mechanism (no RM) are shown as solid lines, while models with a return mechanism (with RM) are shown as dashed lines. It is seen that the return mechanism ensures low event probability with high conditional probability at low *σ_T_*. Lines for the model without a return mechanism end at low *σ_T_* due to insufficient event count (100 events minimum required). Parameters for the return mechanism are *a_X_* = 0.01, *σ_X_* = 0.1, and *β_X_=* 10^−7^. Other parameters are either specified in the figure or standard (see Models, Section II).

We find that the inclusion of a return mechanism leads to enhanced robustness of coupled TA modules. This is indicated by preservation of regular TA module switching dynamics when the toxin RNase activity parameter *β_T_* is sufficiently small (see Fig. 6C,D), which was associated with the system being stuck in the high toxin state without a return mechanism. This increased robustness does not impact our qualitative conclusion that proteolytic coupling leads to strong positive coordination between TA modules. For small *β_T_*, the system with a return mechanism decreases event probabilities while preserving high conditional probabilities (see Fig. 6E), indicating strong positive coordination between events. We conclude that proteolytic coordination can be extended from bistable systems to excitable systems.

## V. CONCLUSIONS

In this article, we examined the role of competition for proteolytic machinery in bacterial persistence. Specifically, we demonstrated that two toxin-antitoxin (TA) modules with antitoxins competing for a common set of proteases robustly display strongly coordinated toxic events. This effect is enhanced but not overshadowed by the coupling of TA modules due to slowed cell growth rate consequent of toxin activity. To demonstrate the influence of proteolytic coupling, we leveraged a novel model motivated by experimentally known molecular species and interactions, and we found using extensive stochastic simulation that such models were tightly coordinated when sharing a common proteolytic pathway. Finally, our findings were extended to a model where TA modules were made more robust by introducing a return mechanism that promotes the reset of the system to a normal physiological state in a reasonable time, as is needed if cells are to return to normal growth after persistence. While the modules considered in this article are identical and based somewhat specifically on the translational inhibition mechanism and complex stoichiometric ratio characteristic of the *mazEF* module in *E. coli*, proteolytic degradation of antitoxins is a common feature of all known TA modules which may differ in complex formation and deleterious toxic effects [2, 7, 8].

Our results suggest more generally that tight coordination of the many TA modules in living cells could occur in large part due to rapid post-translational proteolytic coupling. A mostly unstudied network of proteolytic crosstalk could then be the primary factor behind particular activation patterns of TA modules, and these activation patterns are likely key to understanding the details for persister cell survival. Perhaps the ability for TA modules to mutually activate one another may be a source of redundancy and thus robustness in the activation of persistence, or perhaps proteolytic coupling tunes the particular persister physiology by selectively coordinating TA modules. It is conceivable then that drugs interfering with proteolytic crosstalk could be a major tool for eliminating persistence by providing avenues to new therapies. For example, treatments might be developed to fully clear *M. tuberculosis* infections, which kill approximately 1.6 million people per year [3, 13] and are known for their ability to generate cells with long and robust persistent states [15, 16]. Alternatively, an enhanced understanding of how proteolytic coupling affects persistence may lead to new industrial applications for synthetic biology. A knowledge of how proteolytic coupling affects persistence is likely essential when considering retention of synthetic circuits, especially since many synthetic systems depend on rapid proteolytic degradation, which in turn may suppress persistence. Engineered bacterial persistence could lead to enhanced survival of cells containing synthetic circuits of interest, e.g. in bioreactors or biosensors, leading to reduced cost to maintain these synthetic systems.

## VI. ACKNOWLEDGEMENTS

We thank Nicholas Butzin and Christian Ray for stimulating discussions concerning this work. Funding for this research was provided by the National Science Foundation under Grant No. MCB-1330180.

